# Asymmetry and Heterogeneity in the Plasma Membrane

**DOI:** 10.1101/2025.03.03.641122

**Authors:** Teppei Yamada, Wataru Shinoda

**Author notes:** **Corresponding Author Wataru Shinoda** – Research Institute for Interdisciplinary Science, Okayama University, Okayama 700-8530, Japan.

## Abstract

Plasma membranes (PMs) exhibit asymmetry between their two leaflets in terms of phospholipid headgroups, unsaturation, and resulting membrane properties such as packing and fluidity. Lateral heterogeneity, including the formation of lipid domains, is another crucial aspect of PMs with significant biological implications. However, the nature and even the existence of lipid domains in the two leaflets of PMs remain elusive, hindering a complete understanding of the significance of lipid asymmetry. Using coarse-grained molecular dynamics simulation of the asymmetric PM, we find that the outer leaflet predominantly adopts a liquid-ordered state, whereas the inner leaflet separates into nanoscale (≈10 nm) liquid-ordered and liquid-disordered domains, exhibiting highly dynamic fusion and fission events. This structural asymmetry is further reinforced by asymmetric lateral stress resulting from a cholesterol bias toward the outer leaflet. These findings suggest distinct functional roles for the two leaflets, modulated by asymmetric lateral stress. Additionally, comparing the phase behavior of asymmetric and fully scrambled PMs reveals a key determinant of domain size: intact PMs maintain nanoscale domains, while cell-derived giant PM vesicles, which have lost the strict lipid asymmetry, exhibit microscale domains.

## INTRODUCTION

Cellular plasma membranes (PMs) are complex assemblies composed of a diverse array of lipids and proteins. The lipid bilayer forms the fundamental structure of PMs, compartmentalizing the cytoplasm from the extracellular environment. A key feature of PMs is the asymmetric distribution of lipids between the outer (exoplasmic) and inner (cytoplasmic) leaflets.^1,2^ This asymmetry was initially established regarding phospholipid headgroups: sphingolipids are enriched in the outer leaflet, whereas charged and amino phospholipids are predominant in the inner leaflet.^1,3,4^ Furthermore, a recent lipidomic analysis of the human red blood cell (RBC) PM revealed an apparent asymmetry in phospholipid unsaturation, with the inner leaflet being twice as unsaturated as the outer leaflet.^2^ This unsaturation asymmetry is reflected in the distinct physical properties of the two leaflets, with the outer leaflet being more tightly packed and less fluid than the inner leaflet, suggesting their different functional roles.^2,5^

Another crucial aspect of PMs is lateral heterogeneity, specifically forming lipid domains within each leaflet.^6^ The lipid raft hypothesis proposes the existence of functional ordered domains enriched in cholesterol (CHOL) and sphingolipids within PMs.^7,8^ This hypothesis was initially supported by the isolation of detergent-resistant membranes (DRMs) from PMs^9^ and later corroborated by direct observations of raft-like liquid-ordered (L_o_) phases in synthetic multi-component membranes.^10,11^ However, DRMs are unlikely to accurately reflect native PM structures,^6^ and synthetic membranes lack the lipid complexity of PMs. Giant PM vesicles (GPMVs), derived from intact PMs, exhibit microscale phase separation at near-room temperatures.^12,13^ While GPMVs retain the lipid diversity of PMs, they lose the strict lipid asymmetry.^14^ Consequently, the nature and even the existence of lipid domains in asymmetric PMs remain unclear,^15^ complicating efforts to fully understand the significance of lipid asymmetry.

Molecular dynamics (MD) simulations provide detailed insights into membrane properties and nanoscale lipid organization. In particular, coarse-grained (CG) force fields (FFs), which represent multiple atoms as a single CG bead, enable sufficient sampling to observe lipid domain formation in membranes of biologically relevant size and complexity.^16^ The MARTINI FF is one of the most widely used models for simulating realistic PMs due to its extensive lipid library.^17,18^ While PM models simulated using the MARTINI FF exhibit heterogeneous lipid mixing, they do not show clear phase separation or domain formation.^19,20^ Although this may reflect actual lipid organization in PMs, MARTINI FF has limited accuracy in capturing domain formation within lipid membranes.^21,22^ For instance, no clear phase separation is observed in specific ternary membrane systems, such as those composed of 1,2-dipalmitoylsn-glycero-3-phosphocholine (DPPC), 1,2-dioleoyl-sn-glycero-3-phosphocholine (DOPC), and CHOL, despite experimental evidence of phase separation.^21,22^

SPICA FF is another option for CG MD simulations of lipid membranes.^23–28^ This model was optimized to reproduce key membrane properties, including membrane thickness, area per lipid, and radial distribution functions (RDFs) in binary lipid mixtures, based on all-atom (AA) MD simulations using the CHARMM FF. Recent refinements of the CHOL model in SPICA FF allow accurate descriptions of phase separations in multi-component membranes across various lipid compositions.^28^

In this study, we used CG MD simulations with SPICA FF to investigate the physical properties and lateral heterogeneity of the outer and inner leaflets of the PM. The lipid composition was derived from recent lipidomic data on the human RBC PM.^2^ A CHOL bias toward the outer leaflet induces compressive stress in the outer leaflet and tensile stress in the inner leaflet. This asymmetric lateral stress enhances the differences in membrane physical properties, resulting in a more tightly packed outer leaflet and a more fluid inner leaflet. Order parameter distributions reveal that the outer leaflet is predominantly in the L_o_ phase, while the inner leaflet separates into nanoscale (≈10 nm) L_o_ and liquid-disordered (L_d_) domains, undergoing dynamic fusion and fission events. These findings suggest distinct functional roles for the two leaflets, highlighting the biological significance of lipid asymmetry. Finally, a comparison of phase behavior between asymmetric and fully scrambled PMs identifies a key determinant of domain size: in intact PMs, domains remain at the nanoscale, whereas in GPMVs, which lack the strict lipid asymmetry, domain size expands to the microscale.

## METHODS

### System setup

The initial lipid composition of the simulated PM is shown in Figure 1a. For the phospholipids, we referred to the distilled lipidomes of the human RBC PM.^2^ Lipids with more than 5% content in each leaflet were considered in our simulation. The numbers in the pie chart represent the numbers of carbon atoms and double bonds in two acyl chains for each phospholipid, except for sphingomyelin (SM), for which those of only the acyl chain are shown. In terms of the headgroup of phospholipids, the outer leaflet was composed of PC and SM, while the inner leaflet included phosphatidylethanolamine (PE), PE plasmalogen (PEp), phosphatidylserine (PS), and PC. PEp is the ether form of PE. In addition to this headgroup asymmetry, PM is also largely asymmetric in lipid unsaturation; the inner leaflet is much more unsaturated than the outer one. Here, we call the lipids that include more than four cis double bonds in one tail polyunsaturated lipids. In the outer leaflet, SM lipids were mainly saturated. PC lipids showed relatively low (mono- or di-) unsaturation, except for a polyunsaturated PC (PC 16:0-20:4). Conversely, in the inner leaflet, PE, PEp, and PS lipids were polyunsaturated, with only PC lipids being mono- or diunsaturated. Another notable feature of PM lipidome is an abundance of “hybrid” lipids consisting of one saturated and one unsaturated chain. Phospholipids excluding SM and PE 18:1-20:4 were hybrid lipids. While the CHOL content in each leaflet has been reported in many works, there is no consensus.^29–31^ Some claim that CHOL is abundant in the outer leaflet,^29^ while others claim the opposite.^30^ However, the overall CHOL abundance in RBC is approximately 40 mol%.^2^ Thus, CHOL was included in both leaflets at this concentration in the initial configuration.

**Figure 1.**
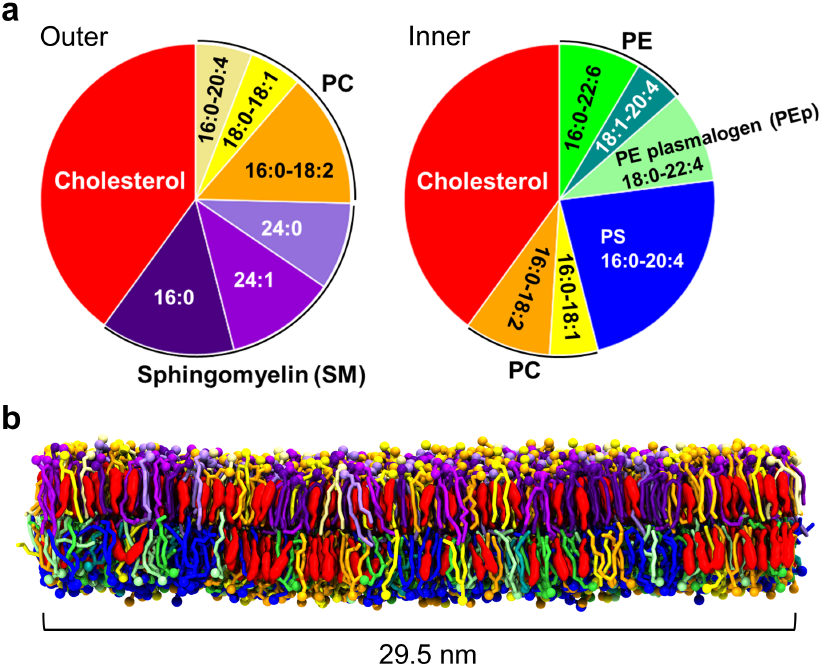
(a) Initial lipid composition of the outer (left) and inner (right) PM leaflets, respectively. (b) Snapshot of the final structure from side view after 120 μs CG MD simulation. Each lipid is shown in the same color as in Figure 1a. The solvent molecules are omitted for clarity.

The number of lipids in the two leaflets is a non-trivial parameter in simulating asymmetric bilayers. Various methods to determine this parameter have been proposed and reviewed recently.^32–35^ Here, we chose the method of matching surface areas. First, we calculated the area of two symmetric bilayers with the lipid compositions of the outer and inner leaflets in Figure 1a. We then set the number of lipids in the two leaflets (outer: 1980 lipids, inner: 1800 lipids) by matching their surface areas to construct the initial configuration. A snapshot of the final structure and detailed system information are shown in Figure 1b and Table S1, respectively.

### Simulation details

We used SPICA FF for all simulations in this study. CG MD simulations were performed using GROMACS version 5.1.5^36^ modified to use SPICA FF. The temperature and pressure were maintained at 298 K and 1 atm, respectively, using the Nosé– Hoover thermostat^37,38^ and the Parrinello–Rahman barostat^39^ with semi-isotropic coupling. Nonbonded LJ interactions were cut off at 1.5 nm, and electrostatic interactions were calculated using the PME method.^40,41^ The time step was set to 10 fs.

### Calculation of the order parameter of phospholipid tails

The order parameter of phospholipid tails was calculated by

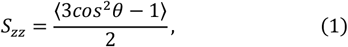

where *θ*is the angle between the bond vectors in the lipid tails and the bilayer normal. We calculated the 2-dimensional distributions of the order parameter defined above for each leaflet to identify lipid domains from the simulation trajectories. The membrane plane was divided into small grids with a length of nm. Phospholipids were assigned to the grids based on their center of mass (COM) positions. We then computed the average order parameter of the phospholipids in the grids. When a grid contained only CHOL and no phospholipids, its order parameter was calculated as the average of the surrounding eight grids. For each time point, the distributions were calculated from 50-ns trajectories.

## RESULTS

### Cholesterol asymmetry induces differential stress

CHOL can flip-flop between the two leaflets over a much shorter time than phospholipids. Only CHOL showed a flip-flop motion in the simulation, while phospholipids did not. To analyze the distribution of CHOL between the two leaflets, we calculated the time variation of the ratio of the number of CHOL in the outer leaflet (*N*_*chol, outer*_) to the total amount of CHOL (*N*_*chol, all*_) (Figure 2a). CHOL molecules were assigned to one leaflet or the other based on the *z*-coordinate of the CG bead corresponding to the hydroxy group. The CHOL distribution was not symmetric (0.5) even at the initial state because CHOL was present in both leaflets at 40 mol%, and the total number of lipids differed between the two leaflets. Figure 2a shows an increase of CHOL in the outer leaflet over time, with about 58% of the total CHOL in the outer leaflet at 120 μs.

**Figure 2.**
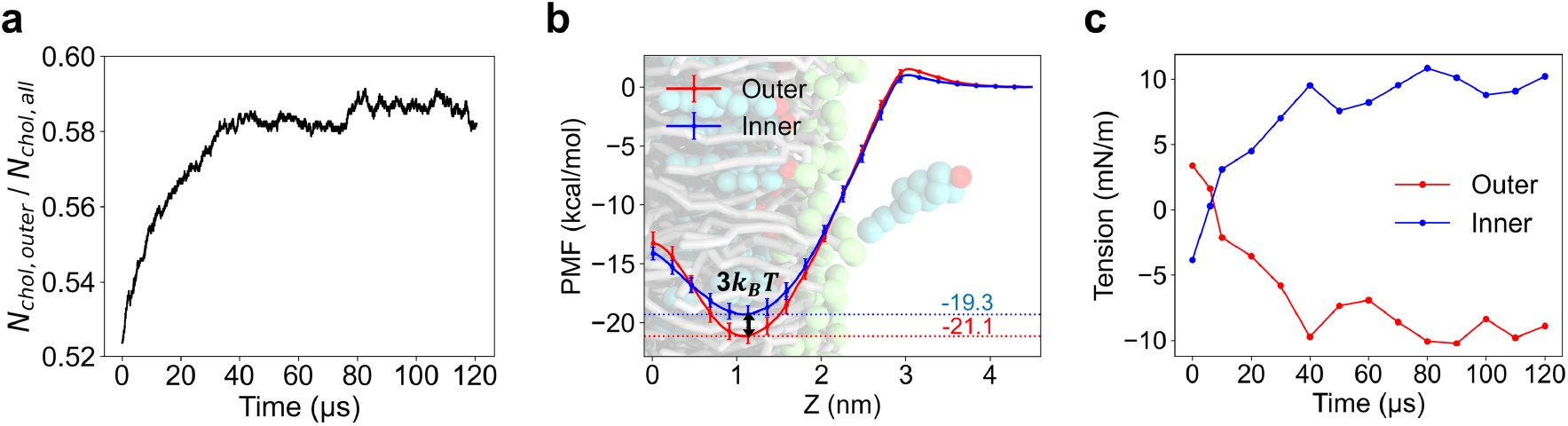
Cholesterol asymmetry induces differential stress. (a) Time variation of the ratio of the number of CHOL in the outer leaflet (*N*_*chol, outer*_) to the total amount of CHOL (*N*_*chol, all*_). (b) Free energy profile of a CHOL along the *z* axis (membrane normal) for the two symmetric bilayers with the lipid composition of the outer and inner leaflets, respectively. The minimum PMF value for each leaflet (outer: −21.1 kcal/mol, inner: −19.3 kcal/mol) and the difference between them (≈−3*k*_*B*_*T*) are specifically noted. (c) Time change of tension acting on each PM leaflet.

CHOL interacts more favorably with SM and saturated lipids than with unsaturated lipids. This difference in affinity would have resulted in an enrichment of CHOL in the SM-rich outer leaflet. To quantify the affinity of CHOL for the two leaflets, we calculated the free energies of CHOL insertion into the two symmetric bilayers with the compositions of the outer and inner leaflets from bulk water (Δ*G*_*outer*_ and Δ*G*_*inner*_) (see the Supporting Information for calculation methods). Figure 2b shows the potential of mean forces (PMFs) of a CHOL along the *z*-axis (bilayer normal). Δ*G*_*outer*_ and Δ*G*_*inner*_ were equal to −21.1 kcal/mol and −19.3 k *G* cal/mol, respectively. Also, the partitioning free energy between the two leaflets, calculated as the difference between Δ*G*_*outer*_ and Δ*G*_*inner*_, was about −3*k*_*B*_*T*, confirming a higher affinity of CHOL for the outer leaflet.

The asymmetric distribution of CHOL resulted in a significant difference in the number of lipids between the two leaflets, suggesting the presence of lateral stress in the membrane. Therefore, we calculated the tension per leaflet from the lateral pressure profile (Figure S1) at each time point (Figure 2c) (see the Supporting Information for calculation methods). The outer leaflet was compressed, while the inner leaflet was under tension. The magnitude of these lateral stresses increased with greater CHOL asymmetry, while the compressive stress in the outer leaflet was nearly canceled out by the tensile stress in the inner leaflet. This occurs because the MD simulation in the NPT ensemble imposed a stress-free condition on the entire bilayer membrane. These asymmetric lateral stresses of equal magnitude have recently been termed as “differential stress”.^42^ The balance among CHOL partitioning free energy, CHOL distribution entropy, and the elastic energy due to differential stress primarily determines transbilayer CHOL distribution in maintained phospholipid asymmetry.^42,43^ Overall, the higher affinity of CHOL for the outer leaflet induced its asymmetric distribution, leading to differential stress acting on both leaflets of the PM.

### Asymmetric physical properties

Here, we analyzed the area per lipid, order parameter, and lateral diffusion coefficient of each leaflet in our PM model, investigating the effects of CHOL asymmetry and the resulting differential stress on these physical properties.

The area per lipid was calculated by dividing the lateral area of the simulation cell by the total number of lipids, including CHOL, in each leaflet (Figure 3a). “Sim-First” represents the average from the initial 1–6 μs of the MD trajectory, while “Sim-Last” denotes the average from the final 5 μs. The lateral stresses at 1–6 μs were sufficiently small compared to the final state (Figure 2c). Thus, comparing Sim-Last with Sim-First reveals the impact of CHOL asymmetry and resulting differential stress on membrane packing.

**Figure 3.**
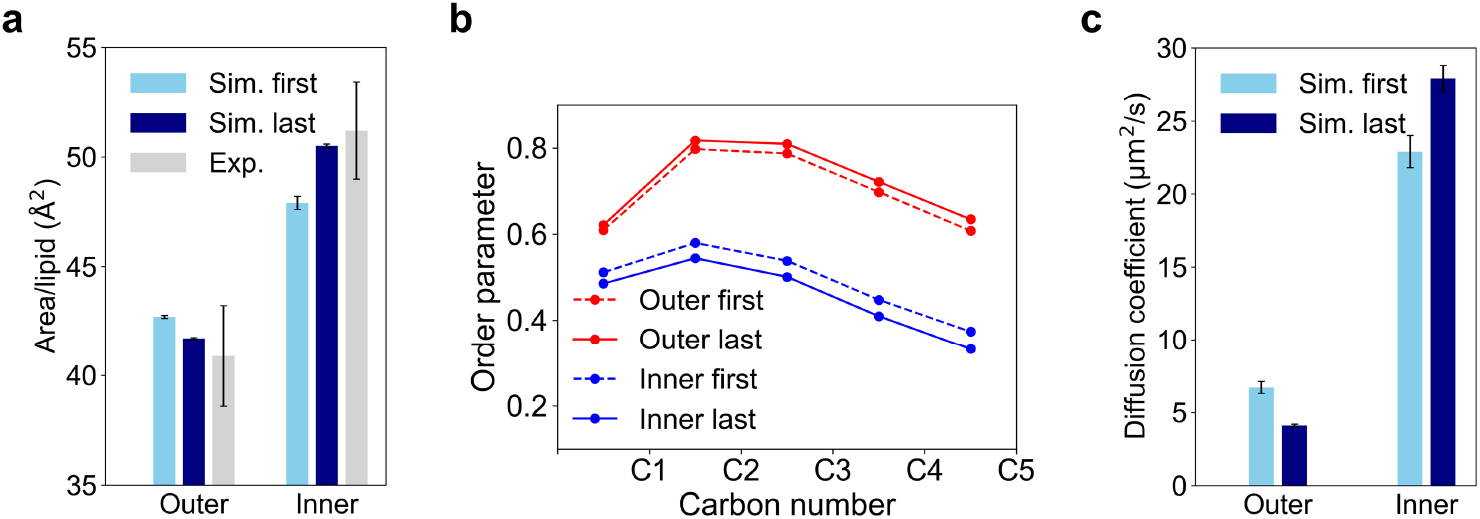
Asymmetric physical properties. (a) Area per lipid of each PM leaflet. “Sim-First” and “Sim-Last” represent the average calculated from the initial 1–6 μs and the final 5 μs MD trajectories, respectively. Experimental data was derived from Reference^44^. (b) Order parameters of phospholipid tails for each PM leaflet. (c) Lateral diffusion coefficients calculated from the average mean square displacement of COM of all lipids for each PM leaflet.

The Sim-Last values indicate that the outer leaflet (41.7 Å^2^/lipid) was more tightly packed than the inner leaflet (50.5 Å^2^/lipid), which is in good agreement with the experimental result for NIH 3T3 fibroblasts (outer: 40.9 Å^2^/lipid, inner: 51.2 Å^2^/lipid).^44^ This packing difference primarily stemmed from the unsaturation asymmetry of lipid acyl chains. Additionally, the packing asymmetry increased from Sim-First to Sim-Last, attributed to differential stress and the “condensing effect”^45^ of CHOL. Compressive stress enhanced the packing of the outer leaflet, while tension reduced that of the inner leaflet. Furthermore, CHOL asymmetry further amplified the packing asymmetry.

The order parameters of phospholipid tails are shown in Figure 3b. The outer leaflet was more ordered than the inner one, with this asymmetry increasing from Sim-First to Sim-Last, similar to the trend observed in the area per lipid.

Finally, we calculated the lateral diffusion coefficients from the average mean square displacements (MSD) of COM of all lipids (Figure S2) for each leaflet (Figure 3c). At the final state, lipid diffusion in the inner leaflet (27.9 μm^2^/s) was approximately seven times faster than in the outer leaflet (4.1 μm^2^/s). The asymmetry in fluidity also increased due to differential stress and the condensing effect of CHOL. Although the diffusion coefficients are not quantitatively accurate due to the use of the CG model, a similar trend has been observed in previous computational studies using all-atom models. In mouse hepatocyte PM, the inner leaflet is approximately five times more fluid than the outer one.^46^ Similarly, in human RBC PM, the inner leaflet is about twice as fluid as the outer one.^2^

### Lateral heterogeneity in the asymmetric plasma membrane

To identify domains in each leaflet, we calculated the two-dimensional distributions of the order parameter. Figure 4a shows the evolution of this distribution in each leaflet during the final 1 μs. The outer leaflet was primarily ordered, except for some small L_d_ domains depleted of CHOL (upper panels in Figure 4a). These L_d_ domains were enriched in the polyunsaturated PC lipid (upper panel in Figure 4b). Examining the positions of the L_d_ domains, the overall structure of the outer leaflet changed little during the 1 μs due to the slow lipid diffusion. In contrast, the inner leaflet separated into CHOL-rich L_o_ domains and CHOL-depleted L_d_ domains (lower panels in Figure 4a). As shown in Figure 4b, the L_o_ domains mainly consisted of CHOL (red) and mono- or di-unsaturated PC lipids (orange), while the L_d_ domains were rich in polyunsaturated lipids (blue). This domain structure dynamically changed, exhibiting reversible fusions and fission events over the 1 μs timeframe due to high fluidity. To emphasize the difference in dynamics between the two leaflets, we focused on the time variation of the order parameter distributions during only final 1 μs. However, the variations over the longer period (final 10 μs) are also provided in Supporting Information.

**Figure 4.**
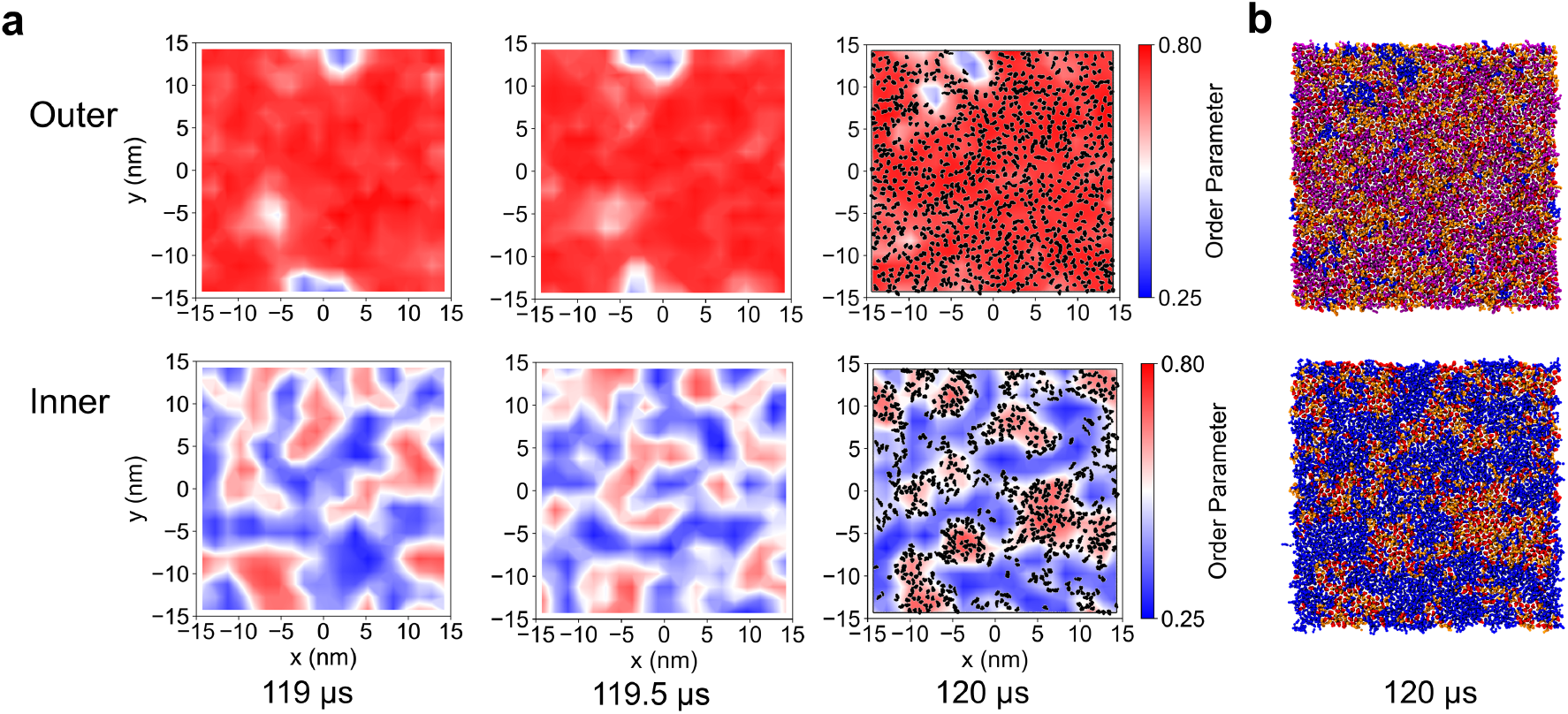
Lateral heterogeneity in the asymmetric PM. (a) Time variation of the two-dimensional order parameter distributions from top view during the final 1 μs for the outer (top) and inner (bottom) leaflets, respectively. CHOL molecules are shown in black at 120 μs. (b) Snapshot of the final structure from top view for each PM leaflet. Color codes are as follows: CHOL (red), SM (purple), mono- or diunsaturated PC (orange), and polyunsaturated lipids (blue).

To further investigate lateral lipid organization, we analyzed the two-dimensional RDF of COM of each lipid around CHOL using the final 1 μs MD trajectories (Figure 5). In the outer leaflet, SM lipids were most likely to be found in the vicinity of CHOL among the phospholipids. Additionally, the first peak of mono- or di-unsaturated PC lipids (orange and yellow) exceeded 1.0, indicating a moderate affinity for CHOL due to their “hybrid” properties. Meanwhile, the polyunsaturated PC (PC 16:0-20:4), which constituted a small fraction of the outer lipidome (9.8 % of phospholipids), exhibited an aversion to CHOL. Thus, most phospholipids in the outer leaflet had an affinity for CHOL. Consequently, CHOL molecules were widely distributed throughout the outer leaflet, contributing to the broadly ordered membrane structure.

**Figure 5.**
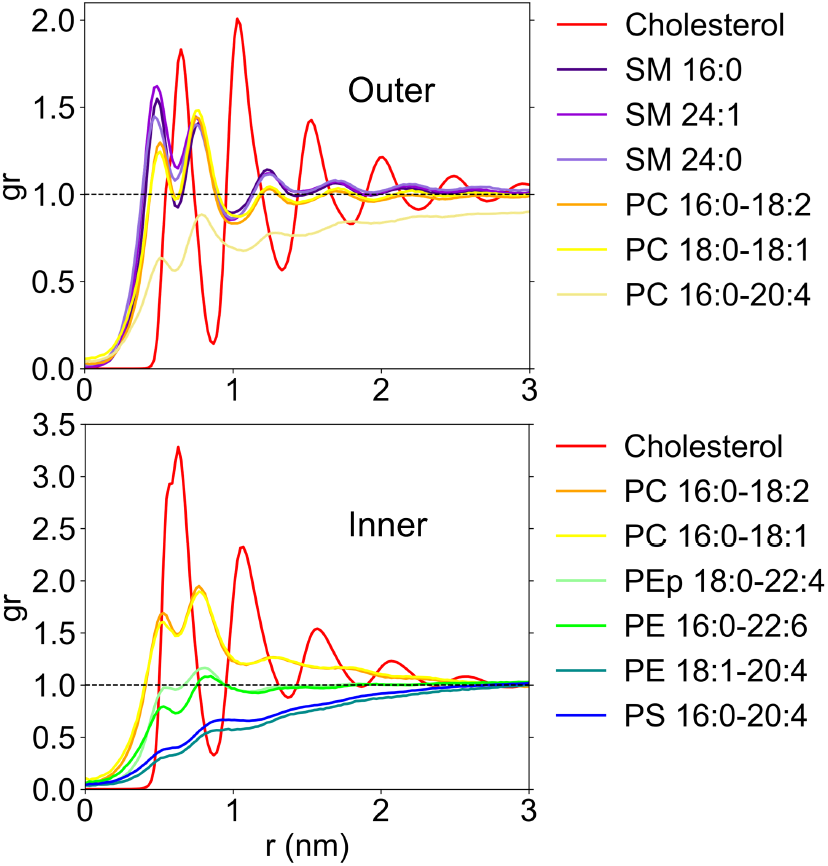
Two-dimensional radial distribution function of COM of each lipid around CHOL for the outer (top) and inner (bottom) PM leaflets, respectively.

In the inner leaflet, mono- or di-unsaturated PC lipids showed a high affinity for CHOL, whereas polyunsaturated PEp, PE, and PS lipids exhibited an aversion to CHOL, as indicated by first peaks below 1.0. This large difference in CHOL affinity among phospholipids led to domain formation in the inner leaflet. Furthermore, the pronounced CHOL-CHOL peaks suggest that L_o_ domains in the inner leaflet were highly enriched in CHOL. Specifically, during the final 1 μs, the average number of CHOL molecules in the inner leaflet was 632, while the total number of mono- or di-unsaturated PC lipids – the primary phospholipid components of L_o_ domains – was 252. Additionally, the strong aversion of PS lipids, which comprised 38.3 % of the total phospholipids in the inner leaflet, to CHOL suggests that PS lipids formed the core of the L_d_ domains, rendering these domains highly negatively charged. This finding aligns with *in vivo* experiments that observed PS clusters in the PM of human embryonic kidney cells.^47^

### Lipid scrambling enlarges lipid domains

The L_o_ domains in the inner leaflet did not grow into a single large domain but remained small (≈10nm). This aligns with the general understanding that lipid domains in PMs are tens of nanometers in size.^15^ In contrast, GPMVs derived from PMs exhibit microscale phase separation near room temperature.^12,13^ The origins of these differing phase behaviors between intact PMs and GPMVs are not fully understood. However, a recent study reported that GPMVs lose lipid asymmetry to some extent (though the precise extent is unknown) and that this loss stabilizes ordered domains.^14^

Here, we aimed to gain deeper molecular insight into these findings and address the question of why domains in intact PMs are nanoscopic. To this end, we performed a simulation on the fully scrambled (symmetric) PM and compared its phase behavior and CHOL-phospholipid interactions with those of the asymmetric PM. The system detail of the scrambled PM is provided in Table S1. Incidentally, the physical properties of the scrambled PM – such as the area per lipid, order parameter, and diffusion coefficient – were approximately the average of the outer and inner leaflets in the asymmetric PM (Figure S3).

Figure 6a shows the two-dimensional order parameter distributions in each leaflet of the scrambled PM. The domains in the scrambled PM were larger than those in the asymmetric PM, which is consistent with experimental observations. As shown in Figure 6b, the L_o_ domains primarily consisted of CHOL (red), SM lipids (purple), and mono- or di-unsaturated PC lipids (orange), while the L_d_ domains were mainly composed of polyunsaturated lipids (blue). Additionally, the domains were anti-registered, meaning that L_d_ domains generally faced L_o_ domains in the opposing leaflet. Several factors contributing to interleaflet coupling have been proposed, including domain thickness mismatch, lipid interdigitation, membrane curvature, and CHOL flip-flop.^48–52^ However, identifying the dominant factors in complex membranes like this remains challenging.

**Figure 6.**
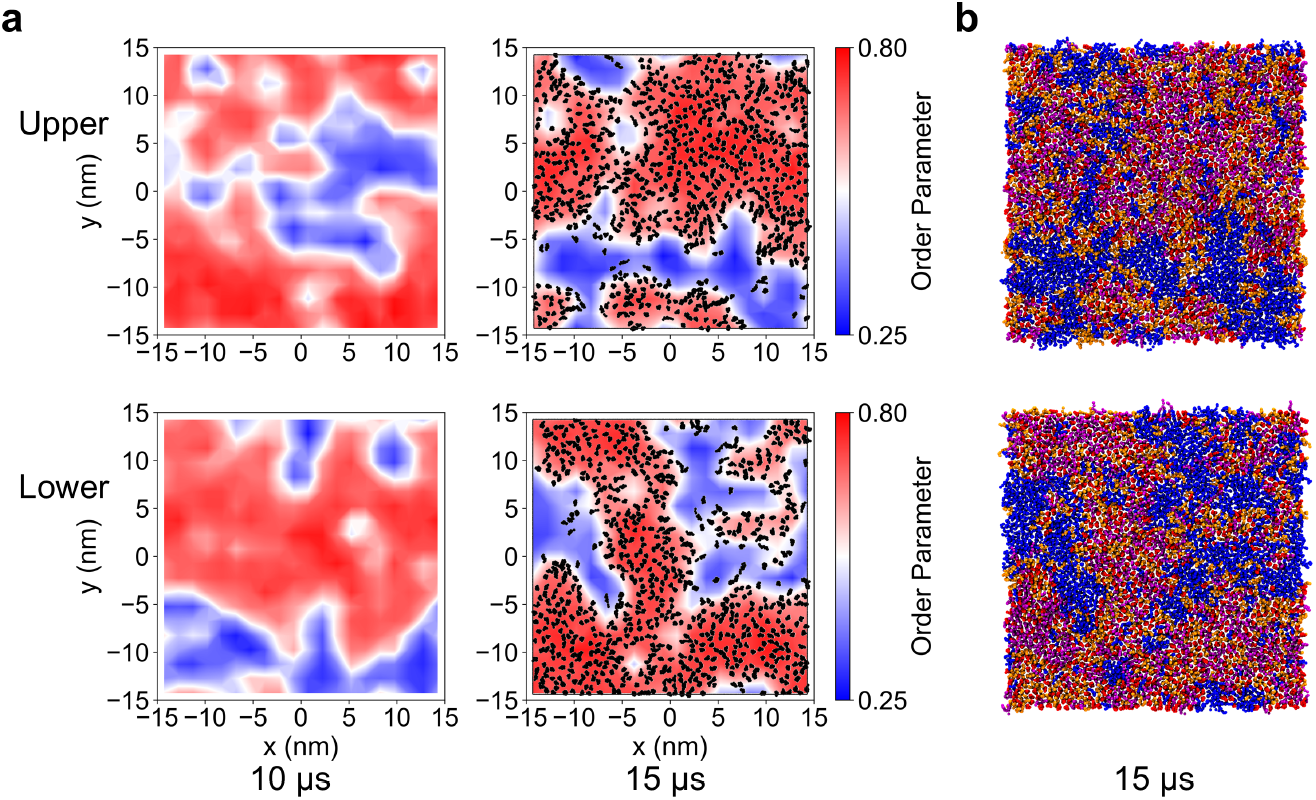
Lateral heterogeneity in the scrambled PM. (a) Two-dimensional order parameter distributions from top view for the upper and lower leaflets in the scrambled PM. CHOL molecules are shown in black at 15 μs. (b) Snapshot of the final structure from top view for each leaflet. The color code is the same as in Figure 4b: CHOL (red), SM (purple), mono- or di-unsaturated PC (orange), and polyunsaturated lipids (blue).

Figure S4 shows the RDF of each lipid around CHOL in the scrambled PM. The peaks of CHOL-SM were higher in the scrambled PM than in the asymmetric PM. Furthermore, the RDF values for CHOL-SM remained above 1.0 even at long distances (>5nm), whereas in the asymmetric PM, they approached 1.0 at around 1.5 nm. These findings indicate that lipid scrambling enhanced the effective interactions between CHOL and SM.

Next, based on the revealed composition of the lipid domains, we discuss the factors that determine domain size in asymmetric and scrambled PMs. A key factor is the differential affinity for CHOL among phospholipids in L_o_ and L_d_ domains. The greater difference in affinity leads to larger domains. Lipid scrambling resulted in the coexistence of SM and a significant amount of polyunsaturated lipids within the same leaflet, a condition absent in the asymmetric PM. SM has a much higher affinity for CHOL compared to polyunsaturated lipids. This strong affinity outweighs mixing entropy, promoting the formation of larger ordered domains. In the inner leaflet of the asymmetric PM, the mono- or di-unsaturated PC lipids, which are the main phospholipid components of L_o_ domains, exhibit only a moderate affinity for CHOL. While this affinity is sufficient to induce nanoscale-ordered domains, it does not drive microscale phase separation.

## DISCUSSION

The lipid raft hypothesis suggests the existence of ordered domains enriched in CHOL, SM, and saturated lipids. However, in our simulation, clear ordered domain formations – specifically, the notable accumulation of CHOL – occurred only in the inner leaflet, which lacked both SM and saturated lipids. The inner leaflet separated into L_o_ domains enriched in CHOL and mono- or di-unsaturated PC lipids and L_d_ domains enriched in polyunsaturated lipids. Does this also occur in actual PMs? Here, we provide support for our findings.

A similar domain formation was recently observed in ternary model membranes composed of CHOL, 1-palmitoyl-2-oleoyl-sn-glycero-3-PC (POPC), and polyunsaturated PE lipids.^53^ This composition is similar to that of the inner leaflet in PMs. Additionally, a key function of lipid domains is the selective recruitment of membrane proteins. Palmitoylation, a post-translational modification, increases the affinity of membrane proteins for ordered domains^54,55^ and primarily occurs in their cytoplasmic regions.^56^ This highlights the importance of inner leaflet domains in selectively recruiting membrane proteins.

Our analysis of the physical properties of the asymmetric PM revealed that the outer leaflet was more tightly packed, more ordered, and less fluid than the inner leaflet. This asymmetry was further amplified by differential stress resulting from CHOL bias toward the outer leaflet. The order parameter distributions showed a widely ordered outer leaflet and coexisting L_o_ and L_d_ domains in the inner leaflet. Based on these findings, we propose distinct functional roles for the two leaflets in PMs. The outer leaflet serves as a barrier against the extracellular environment. Its tightly packed and ordered structure creates a large free energy barrier, restricting molecular passage through the membrane. Meanwhile, the dynamic domains in the inner leaflet laterally sort proteins involved in signal transduction, a process previously associated with lipid rafts.^57^ Non-transmembrane proteins anchored to the inner leaflet via acyl chains – such as Src family kinases, G proteins, and Ras – would particularly benefit from dynamic domain organizations. These proteins can rapidly relocate within the membrane through fast diffusion, facilitating efficient signal transduction. Furthermore, differential stress increased the packing of the outer leaflet while enhancing the fluidity of the inner leaflet, suggesting that it optimizes the functional efficiency of both leaflets.

By comparing the phase behavior of the asymmetric and scrambled PMs, we identified the mechanism by which the loss of lipid asymmetry promotes the formation of larger domains. As mentioned earlier, GPMVs lose strict lipid asymmetry, as evidenced by the exposure of PS lipids on the outer leaflet.^14^ Another approach to study raft domains is DRM isolation from PMs. However, detergents can also induce lipid flip-flop, leading to a loss of lipid asymmetry.^14,58,59^ Thus, DRMs likely reflect larger ordered domains that would not form in intact PMs. Our findings provide valuable insight into the differences in phase behavior between these experimental systems and intact PMs. Furthermore, cells regulate lipid distribution between leaflets, and transient lipid scrambling plays key roles in biological processes such as intercellular communication and intracellular signaling.^60^ Our results offer new insights into these phenomena, contributing to a deeper understanding of membrane organization and function.

In summary, we conducted long-time CG MD simulations of the asymmetric PM using recently reported lipidomic data and analyzed transbilayer CHOL distribution, differential stress, asymmetric physical properties, and lateral heterogeneity. By integrating these findings, we propose distinct functional roles for the two leaflets in PMs, highlighting the significance of lipid asymmetry. Furthermore, comparing the phase behavior of the asymmetric and fully scrambled PMs reveals the molecular determinants of domain size in these systems, providing new insights into the study of GPMVs, DRMs, and the cellular functions associated with transient lipid scrambling.

## ASSOCIATED CONTENT

### Data Availability Statement

The parameters and definitions of CG mapping of SPICA FF are available on the Web page (https://www.spica-ff.org/)” www.spica-ff.org/). The setup tools and GROMACS modified for SPICA FF are available on GitHub (https://github.com/SPICA-group).

### Supporting Information

The Supporting Information is available free of charge.

## AUTHOR INFORMATION

### Authors

**Teppei Yamada** – *Graduate School of Environmental, Life, Natural Science and Technology, Okayama University, Okayama 700-8530, Japan*.

### Notes

The authors declare no competing financial interest.

## ACKNOWLEDGMENT

This work was supported by JST SPRING, Japan Grant Number JPMJSP2126, and JSPS KAKENHI Grant Numbers JP21H01880 and JP24H00843. Calculations were performed on the supercomputer facilities of the Institute for Solid State Physics, the University of Tokyo, and the Research Center for Computational Science, Okazaki, Japan (Project:24-IMS-C090, 23-IMS-C095).

## ABBREVIATIONS

PM: plasma membrane
RBC: red blood cell
DRM: detergent resistant membrane
L_o_: liquid-ordered
L_d_: liquid-disordered
GPMV: giant plasma membrane vesicle
CHOL: cholesterol
DPPC: 1,2-dipalmitoyl-sn-glycero-3-phosphocholine
DOPC: 1,2-dioleoyl-sn-glycero-3-phosphocholine
POPC: 1-palmitoyl-2-oleoyl-sn-glycero-3-phosphocholine
SM: sphingomyelin
PE: phosphatidylethanolamine
PEp: PE plasmalogen
PS: phosphatidylserine
COM: center of mass
PMF: potential of mean force
MSD: mean square displacement
RDF: radial distribution function

---

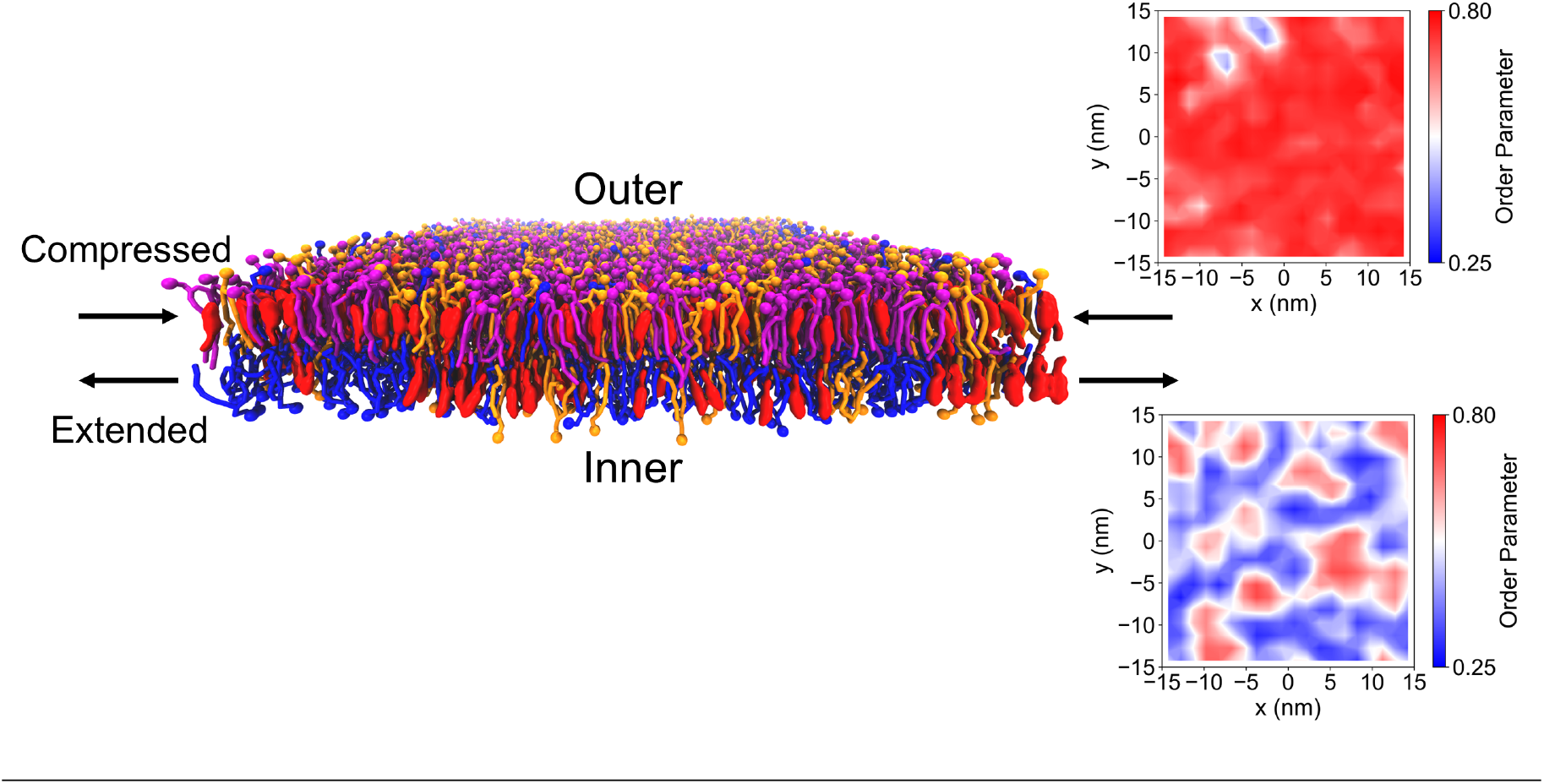

## REFERENCES

(1) Op den Kamp, J. A. Lipid Asymmetry in Membranes. Annu. Rev. Biochem. 1979, 48, 47–71.

(2) Lorent, J. H.; Levental, K. R.; Ganesan, L.; Rivera-Longsworth, G.; Sezgin, E.; Doktorova, M.; Lyman, E.; Levental, I. Plasma Membranes Are Asymmetric in Lipid Unsaturation, Packing and Protein Shape. Nat. Chem. Biol. 2020, 16 (6), 644–652.

(3) Verkleij, A. The Asymmetric Distribution of Phospholipids in the Human Red Cell Membrane. A Combined Study Using Phospholipases and Freeze-Etch Electron Microscopy. Biochim. Biophys. Acta 1973, 323, 178–193.

(4) PK Schick, K. K. G. C. Location of Phosphatidylethanolamine and Phosphatidylserine in the Human Platelet Plasma Membrane. J. Clin. Investig. 1976, 57, 1221–1226.

(5) Gupta, A.; Korte, T.; Herrmann, A.; Wohland, T. Plasma Membrane Asymmetry of Lipid Organization: Fluorescence Lifetime Microscopy and Correlation Spectroscopy Analysis. J. Lipid Res. 2020, 61 (2), 252–266.

(6) Sezgin, E.; Levental, I.; Mayor, S.; Eggeling, C. The Mystery of Membrane Organization: Composition, Regulation and Roles of Lipid Rafts. Nat. Rev. Mol. Cell Biol. 2017, 18 (6), 361–374.

(7) Lingwood, D.; Simons, K. Lipid Rafts as a Membrane-Organizing Principle. Science 2010, 327 (5961), 46–50.

(8) Kusumi, A.; Fujiwara, T. K.; Tsunoyama, T. A.; Kasai, R. S.; Liu, A. A.; Hirosawa, K. M.; Kinoshita, M.; Matsumori, N.; Komura, N.; Ando, H.; Suzuki, K. G. N. Defining Raft Domains in the Plasma Membrane. Traffic 2020, 21 (1), 106–137.

(9) Brown, D. A.; Rose, J. K. Sorting of GPI-Anchored Proteins to Glycolipid-Enriched Membrane Subdomains during Transport to the Apical Cell Surface. Cell 1992, 68 (3), 533–544.

(10) Veatch, S. L.; Keller, S. L. Separation of Liquid Phases in Giant Vesicles of Ternary Mixtures of Phospholipids and Cholesterol. Biophys. J. 2003, 85 (5), 3074–3083.

(11) Veatch, S. L.; Keller, S. L. Miscibility Phase Diagrams of Giant Vesicles Containing Sphingomyelin. Phys. Rev. Lett. 2005, 94 (14), 148101.

(12) Baumgart, T.; Hammond, A. T.; Sengupta, P.; Hess, S. T.; Holowka, D. A.; Baird, B. A.; Webb, W. W. Large-Scale Fluid/Fluid Phase Separation of Proteins and Lipids in Giant Plasma Membrane Vesicles. Proc. Natl. Acad. Sci. U. S. A. 2007, 104 (9), 3165–3170.

(13) Kaiser, H. J.; Lingwood, D.; Levental, I.; Sampaio, J. L.; Kalvodova, L.; Rajendran, L.; Simons, K. Order of Lipid Phases in Model and Plasma Membranes. Proc. Natl. Acad. Sci. U. S. A. 2009, 106 (39), 16645–16650.

(14) Kakuda, S.; Suresh, P.; Li, G.; London, E. Loss of Plasma Membrane Lipid Asymmetry Can Induce Ordered Domain (Raft) Formation. J. Lipid Res. 2022, 63 (1), 100155.

(15) Levental, I.; Levental, K. R.; Heberle, F. A. Lipid Rafts: Controversies Resolved, Mysteries Remain. Trends Cell Biol. 2020, 30 (5), 341–353.

(16) Marrink, S. J.; Corradi, V.; Souza, P. C. T.; Ingólfsson, H. I.; Tieleman, D. P.; Sansom, M. S. P. Computational Modeling of Realistic Cell Membranes. Chem. Rev. 2019, 119 (9), 6184–6226.

(17) Marrink, S. J.; Risselada, H. J.; Yefimov, S.; Tieleman, D. P.; De Vries, A. H. The MARTINI Force Field: Coarse Grained Model for Biomolecular Simulations. J. Phys. Chem. B 2007, 111 (27), 7812–7824.

(18) Souza, P. C. T.; Alessandri, R.; Barnoud, J.; Thallmair, S.; Faustino, I.; Grünewald, F.; Patmanidis, I.; Abdizadeh, H.; Bruininks, B. M. H.; Wassenaar, T. A.; Kroon, P. C.; Melcr, J.; Nieto, V.; Corradi, V.; Khan, H. M.; Domański, J.; Javanainen, M.; Martinez-Seara, H.; Reuter, N.; Best, R. B.; Vattulainen, I.; Monticelli, L.; Periole, X.; Tieleman, D. P.; de Vries, A. H.; Marrink, S. J. Martini3: A General Purpose Force Field for Coarse-Grained Molecular Dynamics. Nat. Methods 2021, 18 (4), 382–388.

(19) Ingólfsson, H. I.; Melo, M. N.; Van Eerden, F. J.; Arnarez, C.; Lopez, C. A.; Wassenaar, T. A.; Periole, X.; De Vries, A. H.; Tieleman, D. P.; Marrink, S. J. Lipid Organization of the Plasma Membrane. J. Am. Chem. Soc. 2014, 136 (41), 14554–14559.

(20) Ingólfsson, H. I.; Carpenter, T. S.; Bhatia, H.; Bremer, P. T.; Marrink, S. J.; Lightstone, F. C. Computational Lipidomics of the Neuronal Plasma Membrane. Biophys. J. 2017, 113 (10), 2271– 2280.

(21) Davis, R. S.; Sunil Kumar, P. B.; Sperotto, M. M.; Laradji, M. Predictions of Phase Separation in Three-Component Lipid Membranes by the MARTINI Force Field. J. Phys. Chem. B 2013, 117 (15), 4072–4080.

(22) Borges-Araújo, L.; Borges-Araújo, A. C.; Ozturk, T. N.; Ramirez-Echemendia, D. P.; Fábián, B.; Carpenter, T. S.; Thallmair, S.; Barnoud, J.; Ingólfsson, H. I.; Hummer, G.; Tieleman, D. P.; Marrink, S. J.; Souza, P. C. T.; Melo, M. N. Martini 3 Coarse-Grained Force Field for Cholesterol. J. Chem. Theory Comput. 2023, 19 (20), 7387–7404.

(23) Shinoda, W.; Devane, R.; Klein, M. L. Multi-Property Fitting and Parameterization of a Coarse Grained Model for Aqueous Surfactants. Mol. Simul. 2007, 33 (1-2), 27–36.

(24) Shinoda, W.; Devane, R.; Klein, M. L. Coarse-Grained Molecular Modeling of Non-Ionic Surfactant Self-Assembly. Soft Matter 2008, 4 (12), 2454–2462.

(25) Shinoda, W.; DeVane, R.; Klein, M. L. Zwitterionic Lipid Assemblies: Molecular Dynamics Studies of Monolayers, Bilayers, and Vesicles Using a New Coarse Grain Force Field. J. Phys. Chem. B 2010, 114 (20), 6836–6849.

(26) Shinoda, W.; Devane, R.; Klein, M. L. Coarse-Grained Force Field for Ionic Surfactants. Soft Matter 2011, 7 (13), 6178–6186.

(27) Shinoda, W.; DeVane, R.; Klein, M. L. Computer Simulation Studies of Self-Assembling Macromolecules. Curr. Opin. Struct. Biol. 2012, 22 (2), 175–186.

(28) Seo, S.; Shinoda, W. SPICA Force Field for Lipid Membranes: Domain Formation Induced by Cholesterol. J. Chem. Theory Comput. 2019, 15 (1), 762–774.

(29) Liu, S. L.; Sheng, R.; Jung, J. H.; Wang, L.; Stec, E.; O’Connor, M. J.; Song, S.; Bikkavilli, R. K.; Winn, R. A.; Lee, D.; Baek, K.; Ueda, K.; Levitan, I.; Kim, K. P.; Cho, W. Orthogonal Lipid Sensors Identify Transbilayer Asymmetry of Plasma Membrane Cholesterol. Nat. Chem. Biol. 2017, 13 (3), 268–274.

(30) Courtney, K. C.; Pezeshkian, W.; Raghupathy, R.; Zhang, C.; Darbyson, A.; Ipsen, J. H.; Ford, D. A.; Khandelia, H.; Presley, J. F.; Zha, X. C24 Sphingolipids Govern the Transbilayer Asymmetry of Cholesterol and Lateral Organization of Model and Live-Cell Plasma Membranes. Cell Rep. 2018, 24 (4), 1037– 1049.

(31) Steck, T. L.; Lange, Y. Transverse Distribution of Plasma Membrane Bilayer Cholesterol: Picking Sides. Traffic 2018, 19 (10), 750–760.

(32) Doktorova, M.; Weinstein, H. Accurate In Silico Modeling of Asymmetric Bilayers Based on Biophysical Principles. Biophys. J. 2018, 115 (9), 1638–1643.

(33) Park, S.; Im, W.; Pastor, R. W. Developing Initial Conditions for Simulations of Asymmetric Membranes: A Practical Recommendation. Biophys. J. 2021, 120 (22), 5041–5059.

(34) Girard, M.; Bereau, T. Induced Asymmetries in Membranes. Biophys. J. 2023, 122 (11), 2092–2098.

(35) Chaisson, E. H.; Heberle, F. A.; Doktorova, M. Building Asymmetric Lipid Bilayers for Molecular Dynamics Simulations: What Methods Exist and How to Choose One? Membranes 2023, 13 (7), 629.

(36) Abraham, M. J.; Murtola, T.; Schulz, R.; Páll, S.; Smith, J. C.; Hess, B.; Lindah, E. GROMACS: High Performance Molecular Simulations through Multi-Level Parallelism from Laptops to Supercomputers. SoftwareX 2015, 1–2, 19–25.

(37) Nosé, S. A Unified Formulation of the Constant Temperature Molecular Dynamics Methods. J. Chem. Phys. 1984, 81 (1), 511– 519.

(38) Hoover, W. G. Canonical Dynamics: Equilibrium Phase-Space Distributions. Phys. Rev. A 1985, 31 (3), 1695–1697.

(39) Parrinello, M.; Rahman, A. Polymorphic Transitions in Single Crystals: A New Molecular Dynamics Method. J. Appl. Phys. 1981, 52 (12), 7182–7190.

(40) Darden, T.; York, D.; Pedersen, L. Particle Mesh Ewald: An N ·log(N) Method for Ewald Sums in Large Systems. J. Chem. Phys. 1993, 98 (12), 10089–10092.

(41) Essmann, U.; Perera, L.; Berkowitz, M. L.; Darden, T.; Lee, H.; Pedersen, L. G. A Smooth Particle Mesh Ewald Method. J. Chem. Phys. 1995, 103 (19), 8577–8593.

(42) Hossein, A.; Deserno, M. Spontaneous Curvature, Differential Stress, and Bending Modulus of Asymmetric Lipid Membranes. Biophys. J. 2020, 118 (3), 624–642.

(43) Varma, M.; Deserno, M. Distribution of Cholesterol in Asymmetric Membranes Driven by Composition and Differential Stress. Biophys. J. 2022, 121 (20), 4001–4018.

(44) Doktorova, M.; Symons, J. L.; Zhang, X.; Wang, H.-Y.; Schlegel, J.; Lorent, J. H.; Heberle, F. A.; Sezgin, E.; Lyman, E.; Levental, K. R.; Levental, I. Cell Membranes Sustain Phospholipid Imbalance via Cholesterol Asymmetry. bioRxiv 2024, 2023.07.30.551157.

(45) Hung, W. C.; Lee, M. T.; Chen, F. Y.; Huang, H. W. The Condensing Effect of Cholesterol in Lipid Bilayers. Biophys. J. 2007, 92 (11), 3960–3967.

(46) Andoh, Y.; Hayakawa, S.; Okazaki, S. Molecular Dynamics Study of Lipid Bilayers Modeling Outer and Inner Leaflets of Plasma Membranes of Mouse Hepatocytes. I. Differences in Physicochemical Properties between the Two Leaflets. J. Chem. Phys. 2020, 153 (3).

(47) Bobone, S.; Hilsch, M.; Storm, J.; Dunsing, V.; Herrmann, A.; Chiantia, S. Phosphatidylserine Lateral Organization Influences the Interaction of Influenza Virus Matrix Protein 1 with Lipid Membranes. J. Virol. 2017, 91 (12).

(48) Putzel, G. G.; Uline, M. J.; Szleifer, I.; Schick, M. Interleaflet Coupling and Domain Registry in Phase-Separated Lipid Bilayers. Biophys. J. 2011, 100 (4), 996–1004.

(49) Seo, S.; Murata, M.; Shinoda, W. Pivotal Role of Interdigitation in Interleaflet Interactions: Implications from Molecular Dynamics Simulations. J. Phys. Chem. Lett. 2020, 11 (13), 5171– 5176.

(50) Perlmutter, J. D.; Sachs, J. N. Interleaflet Interaction and Asymmetry in Phase Separated Lipid Bilayers: Molecular Dynamics Simulations. J. Am. Chem. Soc. 2011, 133 (17), 6563– 6577.

(51) Thallmair, S.; Ingólfsson, H. I.; Marrink, S. J. Cholesterol Flip-Flop Impacts Domain Registration in Plasma Membrane Models. J. Phys. Chem. Lett. 2018, 9 (18), 5527–5533.

(52) Fujimoto, T.; Parmryd, I. Interleaflet Coupling, Pinning, and Leaflet Asymmetry-Major Players in Plasma Membrane Nanodomain Formation. Front. Cell Dev. Biol. 2017, 4, 155.

(53) Goh, M. W. S.; Tero, R. Non-Raft Submicron Domain Formation in Cholesterol-Containing Lipid Bilayers Induced by Polyunsaturated Phosphatidylethanolamine. Colloids Surfaces B Biointerfaces 2022, 210, 112235.

(54) Levental, I.; Lingwood, D.; Grzybek, M.; Coskun, Ü.; Simons, K. Palmitoylation Regulates Raft Affinity for the Majority of Integral Raft Proteins. Proc. Natl. Acad. Sci. U. S. A. 2010, 107 (51), 22050–22054.

(55) Lorent, J. H.; Diaz-Rohrer, B.; Lin, X.; Spring, K.; Gorfe, A. A.; Levental, K. R.; Levental, I. Structural Determinants and Functional Consequences of Protein Affinity for Membrane Rafts. Nat. Commun. 2017, 8, 1219.

(56) Jiang, H.; Zhang, X.; Chen, X.; Aramsangtienchai, P.; Tong, Z.; Lin, H. Protein Lipidation: Occurrence, Mechanisms, Biological Functions, and Enabling Technologies. Chem. Rev. 2018, 118 (3), 919–988.

(57) Simons, K.; Toomre, D. Lipid Rafts and Signal Transduction. Nat. Rev. Mol. Cell Biol. 2000, 1 (1), 31–39.

(58) Pantaler, E.; Kamp, D.; Haest, C. W. M. Acceleration of Phospholipid Flip-Flop in the Erythrocyte Membrane by Detergents Differing in Polar Head Group and Alkyl Chain Length. Biochim. Biophys. Acta - Biomembr. 2000, 1509 (1–2), 397–408.

(59) Dietel, L.; Kalie, L.; Heerklotz, H. Lipid Scrambling Induced by Membrane-Active Substances. Biophys. J. 2020, 119 (4), 767– 779.

(60) Doktorova, M.; Symons, J. L.; Levental, I. Structural and Functional Consequences of Reversible Lipid Asymmetry in Living Membranes. Nat. Chem. Biol. 2020, 16 (12), 1321–1330.

