## Supplementary material for "Asymmetry and Heterogeneity in the Plasma Membrane": SI.pdf

**Partitioning free energy of cholesterol between the outer and inner leaflets**

To calculate the partitioning free energy of cholesterol (CHOL) between the outer and inner leaflets, we calculated the free energy of CHOL insertion into the two symmetric bilayers, each corresponding to the composition of the outer ($\Delta G_{outer}$) and inner ($\Delta G_{inner}$) leaflets, respectively. The partitioning free energy was then obtained as the difference between $\Delta G_{outer}$ and $\Delta G_{inner}$. We built small symmetric bilayers for potential mean force (PMF) calculations, with the outer bilayer containing 440 lipids and the inner bilayer containing 400 lipids. PMFs were calculated using umbrella sampling (US)[1] with the weighted histogram analysis method (WHAM)[2]. The reaction coordinate was defined as the projected distance to the bilayer normal ($z$-direction) between the center of mass (COM) of a CHOL and the COM of the lipid bilayer. Along the reaction coordinate ($z$), 23 US windows were used, centered at $z = 0$ nm, 0.2 nm, …, 4.4 nm, with a force constant of 100 kcal $nm^{-2}$ $mol^{-1}$ for each window. Each window underwent 1 μs of sampling simulations. The averages and standard deviations of the PMFs were estimated from the block data obtained from every 200 ns of sampling.

**Leaflet tension.**

The tension per the outer and inner leaflets ($T_{out}$ and $T_{inn}$) were calculated using the following equations.

$$T_{out} = -\int_{0}^{L_z/2} P(z)\, dz, \quad (1)$$

$$T_{inn} = -\int_{-L_z/2}^{0} P(z)\, dz, \quad (2)$$

$$P(z) = \frac{1}{2}\left[P_{xx}(z) + P_{yy}(z)\right] - P_{zz}(z), \quad (3)$$

where the $P_{ii}(z)$ are the diagonal components of the local pressure tensor in each slab at position $z$, and $L_z$ is the length of the simulation box in the direction of the $z$-axis. The local pressure tensors were calculated from 1 ns trajectory (1000 frame) for each time using GROMACS-LS,[3,4] a modified version.

**Table 1.** System information of the simulated PM models.

| | outer (upper) leaflet | | inner (lower) leaflet | | solvent (0.15 M NaCl) | |
|---|---|---|---|---|---|---|
| | **lipid** | **number** | **lipid** | **number** | **water/ions** | **number** |
| **asymmetric PM** | CHOL | 792 | CHOL | 720 | Water | 140657 |
| | SM 16:0 | 279 | PS 16:0-20:4 | 414 | $Na^+$ | 787 |
| | SM 24:1 | 225 | PEp 18:0-22:4 | 171 | $Cl^-$ | 373 |
| | SM 24:0 | 180 | PE 16:0-22:6 | 153 | | |
| | PC 16:0-18:2 | 279 | PE 18:1-20:4 | 90 | | |
| | PC 18:0-18:1 | 108 | PC 16:0-18:2 | 162 | | |
| | PC 16:0-20:4 | 117 | PC 16:0-18:1 | 90 | | |
| **total** | | 1980 | | 1800 | | |
| **symmetric (outer, small)** | CHOL | 88 | Same as in the upper leaflet. | | Water | 23084 |
| | SM 16:0 | 31 | | | $Na^+$ | 61 |
| | SM 24:1 | 25 | | | $Cl^-$ | 61 |
| | SM 24:0 | 20 | | | | |
| | PC 16:0-18:2 | 31 | | | | |
| | PC 18:0-18:1 | 12 | | | | |
| | PC 16:0-20:4 | 13 | | | | |
| **total** | | 220 | | | | |
| **symmetric (inner, small)** | CHOL | 80 | Same as in the upper leaflet. | | Water | 22792 |
| | PS 16:0-20:4 | 46 | | | $Na^+$ | 153 |
| | PEp 18:0-22:4 | 19 | | | $Cl^-$ | 61 |
| | PE 16:0-22:6 | 17 | | | | |
| | PE 18:1-20:4 | 10 | | | | |
| | PC 16:0-18:2 | 18 | | | | |
| | PC 16:0-18:1 | 10 | | | | |
| **total** | | 200 | | | | |
| **scrambled PM** | CHOL | 756 | Same as in the upper leaflet. | | Water | 143838 |
| | SM 16:0 | 135 | | | $Na^+$ | 783 |
| | SM 24:1 | 117 | | | $Cl^-$ | 369 |
| | SM 24:0 | 90 | | | | |
| | PC 16:0-18:2 | 225 | | | | |
| | PC 16:0-18:1 | 45 | | | | |
| | PC 18:0-18:1 | 54 | | | | |
| | PC 16:0-20:4 | 54 | | | | |
| | PS 16:0-20:4 | 207 | | | | |
| | PEp 18:0-22:4 | 90 | | | | |
| | PE 16:0-22:6 | 72 | | | | |
| | PE 18:1-20:4 | 45 | | | | |
| **total** | | 1890 | | | | |

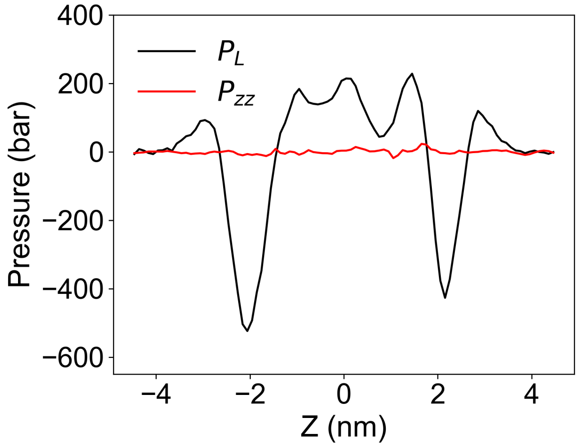


**Figure S1.** Pressure profile along *z*-axis (membrane normal) in the asymmetric PM at 120 μs. The origin corresponds to the center of the bilayer. The region where *z* is greater than 0 corresponds to the outer leaflet, while the region where *z* is less than 0 corresponds to the inner leaflet. $P_L$ represents the average of $P_{xx}$ and $P_{yy}$.

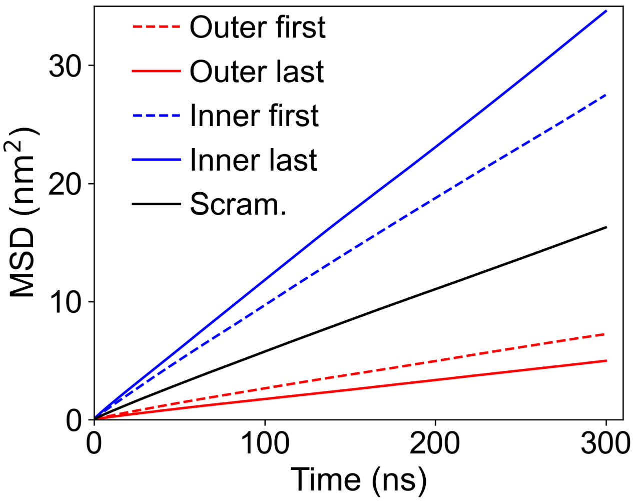


**Figure S2.** Average mean square displacements of COM of all lipids for the asymmetric and scrambled PM. “Outer” and “Inner” refer to the two leaflets in the asymmetric PM. “First” and “Last” represent the values calculated from the initial 1-2 μs and the final 1 μs MD trajectories, respectively. “Scram.” indicates the value calculated from the final 1 μs MD trajectories of the scrambled PM.

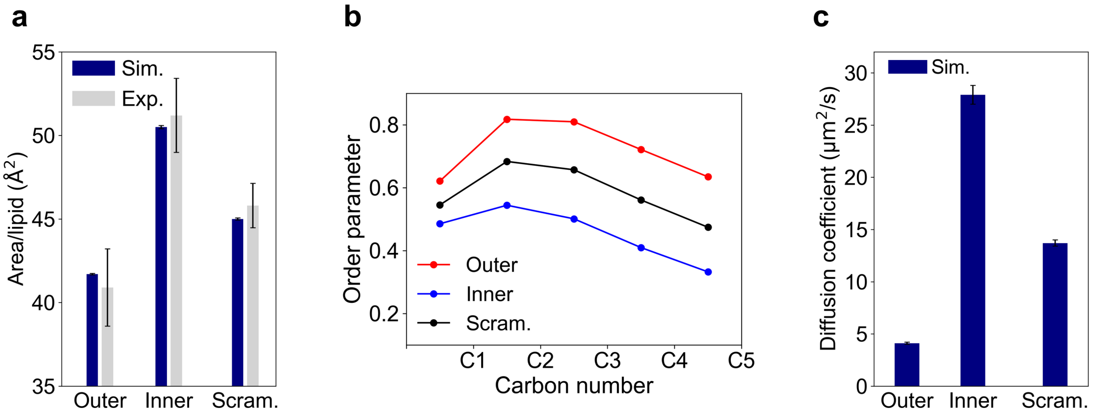


**Figure S3.** Comparison of physical properties between the asymmetric and scrambled PM. (a) Area per lipid calculated for the two leaflets in the asymmetric and scrambled PM. The values represent averages obtained from the final 5 μs of MD trajectories. Experimental data were derived from Reference[5]. (b) Order parameters of phospholipid tails for the two leaflets in the asymmetric and scrambled PM. (c) Lateral diffusion coefficients calculated from the average mean square displacement of COM of all lipids for the two leaflets in the asymmetric and scrambled PM.

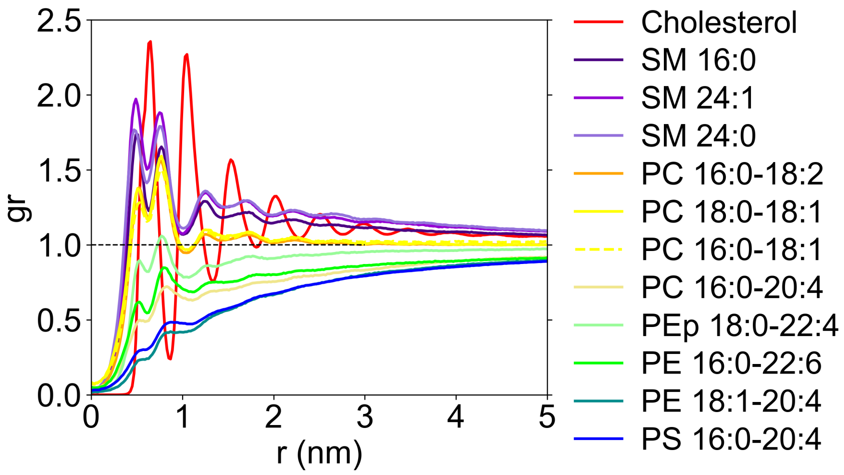


**Figure S4.** Two-dimensional radial distribution function of COM of each lipid around CHOL in the scrambled PM, calculated from the final 1 μs of MD trajectories.

### References


(1) Torrie, G. M.; Valleau, J. P. Nonphysical Sampling Distributions in Monte Carlo Free-Energy Estimation: Umbrella Sampling. *J. Comput. Phys.* **1977**, *23* (2), 187–199.

(2) Kumar, S.; Rosenberg, J. M.; Bouzida, D.; Swendsen, R. H.; Kollman, P. A. Multidimensional Free-Energy Calculations Using the Weighted Histogram Analysis Method. *J. Comput. Chem.* **1995**, *16* (11), 1339–1350.

(3) Ollila, O. H. S.; Risselada, H. J.; Louhivuori, M.; Lindahl, E.; Vattulainen, I.; Marrink, S. J. 3D Pressure Field in Lipid Membranes and Membrane-Protein Complexes. *Phys. Rev. Lett.* **2009**, *102* (7), 078101.

(4) Vanegas, J. M.; Torres-Sánchez, A.; Arroyo, M. Importance of Force Decomposition for Local Stress Calculations in Biomembrane Molecular Simulations. *J. Chem. Theory Comput.* **2014**, *10* (2), 691–702.

(5) Doktorova, M.; Symons, J. L.; Zhang, X.; Wang, H.-Y.; Schlegel, J.; Lorent, J. H.; Heberle, F. A.; Sezgin, E.; Lyman, E.; Levental, K. R.; Levental, I. Cell Membranes Sustain Phospholipid Imbalance via Cholesterol Asymmetry. *bioRxiv* **2024**, 2023.07.30.551157.
